# Morphological and molecular effects of short-term water deficiency in barley stamen maturation

**DOI:** 10.1101/2024.11.19.624362

**Authors:** Richard Lange, Youjun Zhang, Alisdair R. Fernie, Iván F. Acosta

## Abstract

Water deficiency at the reproductive stage of cereal crops mainly affects the development or function of male organs, which causes strong losses in grain yield. We investigated the effects of short-term drought on the post-meiotic maturation of stamens in barley cultivar Scarlett, where it leads to a stage-dependent decline in fertility. Water deficiency neither affects pollen viability nor the formation of trinuclear pollen. However, it completely blocks pollen starch accumulation. Metabolite profiling suggests that this is due to decreased sugar content at starch-filling stages, probably reflecting impaired carbon supply from photosynthetic tissues to anther sinks. Accordingly, transcriptomic analysis shows that drought reduces the expression of stamen sugar transporters. Moreover, drought causes a strong downregulation of the pollen transcriptional network of auxin signalling and central carbon metabolism genes that controls barley pollen starch production. This wider model of the molecular effects of water deficiency on cereal pollen provides a solid foundation to characterize tolerance mechanisms in potential drought-resistant germplasm.

## INTRODUCTION

Throughout their life cycle, plants are exposed to a range of unfavourable environmental conditions. Among them, drought stress is a major threat that jeopardizes rain-fed agricultural crop production by limiting the vegetative and reproductive capability of plants (Barnabas et al., 2008). Between 1983 and 2009, drought affected approximately 75% of the global harvested areas of crops, causing economic losses of 166 billion U.S. dollars (Kim et al., 2019). Driven by anthropogenic-caused climate change, the adverse impact of drought and heatwaves on crop production has tripled throughout the last decades (Brás et al., 2021). In cereals, which provide the majority of consumed calories globally (Poole et al., 2021), droughts are cumulatively and consistently diminishing production, overall imposing 7% more damage to yields than in the past (Lesk et al., 2016; Brás et al., 2021). A recent meta-analysis has estimated that drought stress reduces wheat and rice yields by an average of 27.5% and 25.4%, respectively (Zhang et al., 2018). As climate change continues to alter the global environment, the frequency and intensity of drought periods will likely increase in the future (Hoerling et al., 2012; Dai, 2013; Lehner et al., 2017; Haile et al., 2020; Hari et al., 2020), thus posing an elevated threat to crop yields.

In rice and wheat, cold or drought stresses cause the highest loss of grain number when they occur during the reproductive phase (Oliver et al., 2005; Ji et al., 2010; Jin et al., 2013). The development of male reproductive organs is especially vulnerable to drought (Dorion et al., 1996; Liu and Bennett, 2011; Jin et al., 2013), in contrast to female reproductive structures, which mostly remain unaffected (Bingham, 1966; Saini and Aspinall, 1981; Ji et al., 2010). Therefore, male sterility is widely considered the major driver of grain loss in cereals subjected to drought (Dolferus et al., 2011; Yu et al., 2019).

An important part of male reproductive development in flowering plants is the formation and maturation of male gametophytes in the form of pollen grains (Scott et al., 2004). First, differentiation of sporogenous cells and subsequent meiosis (microsporogenesis) give rise to haploid microspores that are released into the anther locule (Scott et al., 2004; Wilson and Zhang, 2009). Subsequently, microspores acquire a multi-layered pollen wall and become vacuolized (Christensen et al., 1972; Owen and Makaroff, 1995). In cereal crops, developing gametophytes undergo two rounds of asymmetric mitosis to become tricellular, thus containing two generative cells and one vegetative cell at maturity (Eady et al., 1995). Moreover, late stages of pollen development are characterized by the accumulation of large starch stocks that are essential for pollen function (Amanda et al., 2022). Finally, anther sacs open to release mature pollen that goes on to germinate and elongate a pollen tube, which carries the generative cells to the female gametophyte for fertilization.

Research within the last decades has identified meiosis and the stages shortly after microspore release as the most vulnerable phases to drought stress (Dolferus et al., 2011; Yu et al., 2019). For example, water deficit during male meiosis in wheat, rice and tomato can lead to morphological defects and impaired function of the tapetum (Lalonde et al., 1997; Ji et al., 2010; Guo et al., 2013; Jin et al., 2013; Lamin-Samu et al., 2021), and anthers can become smaller, pale and shrivelled (Sheoran and Saini, 1996; Jin et al., 2013). Moreover, histological cross-sections of drought-stressed rice anthers reveal hypertrophy of endothecial cells and deformation of whole locules, suggesting that intense drought can impact all anther tissues (Jin et al., 2013; Guo et al., 2016). Drought-induced male sterility has been frequently linked to abnormal microspore development (Yu et al., 2019), including microspore disorientation, irregular vacuolation and, in extreme cases, complete degeneration (Jin et al., 2013; Guo et al., 2016). Pollen of drought-stressed anthers often show wall defects and reduced or complete absence of starch (Saini et al., 1984; Sheoran and Saini, 1996; Lalonde et al., 1997; Jin et al., 2013).

In cereals, all these drought-induced defects have been associated with impaired carbohydrate metabolism (Saini et al., 1984; Dorion et al., 1996; Sheoran and Saini, 1996; Lalonde et al., 1997; Jin et al., 2013). Short-term drought stress in anthers of wheat and rice irreversibly represses the activity of acid invertases (IVR), important enzymes of sugar utilization (Sheoran and Saini, 1996; Lalonde et al., 1997; Koonjul et al., 2005; Nguyen et al., 2010); and of soluble starch-synthase and ADP-glucose pyrophosphorylase, two key enzymes required for starch biosynthesis (Dorion et al., 1996). Moreover, drought downregulates the expression of genes encoding not only invertases but also glycoside hydrolases, 6-phosphofructokinase and UDP-galactose/UDP-glucose transporters (Koonjul et al., 2005; Nguyen et al., 2010; Jin et al., 2013). It has therefore been proposed that maintenance of anther sink strength through elevated expression of genes participating in carbohydrate metabolism can contribute to drought tolerance (Ji et al., 2010). For example, increased transcript levels in anthers of genes encoding the cell wall invertase OsINV4 and the monosaccharide transporter OsMST8 could partially explain drought and heat tolerance of rice cultivar N22 (Li et al., 2015). Furthermore, expression of the invertase gene *TaIVR1* remained unaffected or increased significantly under drought-stress in drought-tolerant wheat accessions SYN604 and Jinmai47, respectively, whereas it was mostly repressed in drought-susceptible wheat lines (Ji et al., 2010; Dong et al., 2017).

Drought stress also affects hormone homeostasis in anthers (Yu et al., 2019). Abscisic acid (ABA), a central player in the regulation of abiotic stress responses (Sah et al., 2016; Weiste et al., 2017), accumulates in drought-stressed male reproductive tissues and correlates with repression of carbohydrate metabolism genes, thus contributing to pollen sterility (Morgan, 1980; Ji et al., 2010; Liu and Bennett, 2011). Drought imposed during the anthesis stage of rice panicles decreased the endogenous content of indole-3-acetic-acid (IAA), the main bioactive auxin in plants, and downregulated genes encoding auxin synthesis enzymes, auxin co-receptors and auxin response factors (ARFs) (Sharma et al., 2018). Accordingly, exogenous application of IAA mitigated the negative effects of heat and drought stress on pollen viability and overall spikelet fertility (Sharma et al., 2018). Auxin application during pre-anthesis drought stress in Arabidopsis, also improves tolerance to the stress, possibly due to altered regulation of reactive oxygen species (ROS), sugar metabolism and stress-related genes (Shi et al., 2014).

To ensure sufficient grain yield in the near future, it is urgent to improve drought tolerance in cereal crops (Farooq et al., 2009). However, this requires in-depth knowledge of the mechanisms that orchestrate plant responses to the stress and identification of the most vulnerable aspects of development (Farooq et al., 2009; Dolferus et al., 2011). Although adverse effects of pre-anthesis drought stress on grain formation in barley are known (Rajala et al., 2011), detailed descriptions of drought-induced changes during barley stamen maturation are lacking. Thus, in addition to time-resolved morphological studies, we carried out transcriptome and metabolite analyses in maturing barley stamens to obtain broader insights into stage-dependent dynamics of drought responses. The results converge to a model where water deficiency impairs not only photosynthate accumulation in barley stamens but also the transcriptional network that auxin normally activates to enhance sugar utilization and energy production during pollen maturation.

## RESULTS

### Staging the phase of stamen maturation in barley cultivar Scarlett

The barley inflorescence (“spike”) is hidden inside the leafy pseudostem during most of its development. Under our greenhouse conditions, it extrudes out of the sheath of the last (flag) leaf only after fertilization. Thus, destructive dissection is required to determine the exact stages of stamen development in a given inflorescence, which prevents further experimentation. Therefore, we first established a non-destructive method to predict stamen development in hidden Scarlett inflorescences, based on three tiller features: the flag leaf sheath extension out of the penultimate leaf, the distance of the hidden spike tip to the auricle of the penultimate leaf, and the perceived length of the hidden spike (Figure 1A). These features correlated with advancing anther and pistil development, as determined with the Waddington (W) morphological scale that we modified previously (Figure 1B, C) (Amanda et al., 2022). Scarlett anthers at stage W7.5 harbor either tetrads or recently released microspores (Figure 1D), which represent the end of meiosis. At stages W8 and W8.25, the thickening wall and the expansion of free microspores indicate the initiation of stamen and pollen maturation (Figure 1D).

**Figure 1.**
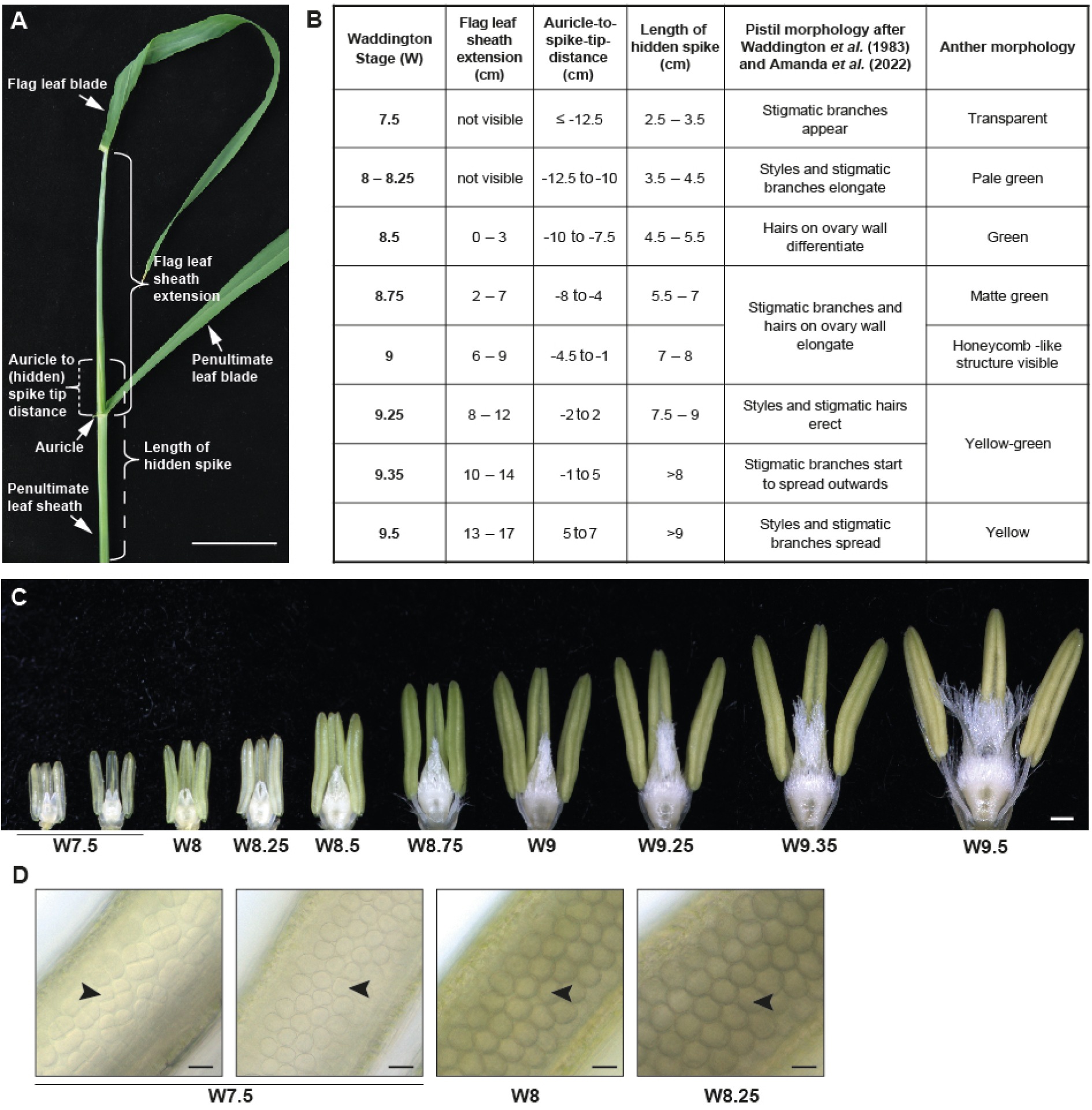
Morphological scale of stamen maturation in cultivar Scarlett under normal greenhouse conditions. **A**) Tiller morphology and features used to predict floral organ development. Bar, 5 cm. **B**) Modified Waddington (W) scale of pistil and anther development and its correlation with external tiller features. The hidden spike tip is positioned below or above the auricle of the penultimate leaf at early or late stages, respectively, resulting in negative or positive auricle-to-spike-tip distances. **C**) Stamen and pistil morphology between stages W7.5 and W9.5. Bar, 500 µm. **D**) Male gametophytes visualized through fresh anthers mounted in perfluorperhydrophenanthren. Arrowheads indicate tetrads (W7.5, left), recently released microspores (W7.5, right) and expanding free microspores with thicker cell walls (W8 and W8.25). Bars, 200 µm.

### Short-term drought stress during stamen maturation affects vegetative and reproductive development

Next, we established a system to perform short-term drought treatments during the stamen maturation phase in pairs of Scarlett plants grown in 1-liter pots under greenhouse conditions. We began experiments approximately 6 weeks after sowing, when the flag leaf sheath of the first tiller started extending and we could tag and predict the stage of stamen development in the first three tillers. Maintaining the soil water content at ∼25% in control pots sustained the relative water content of the penultimate leaf above 90% (Figure 2A). Instead, withholding water dropped the soil water content to around 0% after 2 days and to completely undetectable levels after 3 days, which ultimately reduced the average leaf water content to 57% (Figure 2A). Moreover, drought-stressed plants exhibited leaf wilting and reduced height by the end of the 3-day treatment (Figure 2B). Thereafter, we re-watered these plants, maintained them at control levels and examined spike morphology approximately two weeks later. Control inflorescences appeared mostly well hydrated and with grain-filled spikelets (Figure 2C, D). Although some inflorescences from drought-stressed plants also appeared hydrated, they showed various degrees of empty (sterile) spikelets (Figure 2C). Other inflorescences either carried a fraction of pale and dehydrated spikelets, predominantly at the top of the spike, or were completely dehydrated, thin and stunted (Figure 2C). Inflorescences that were stressed at earlier stages of stamen maturation tended to show more hydrated morphologies, while partial and fully dehydrated inflorescences tended to occur more often when we started the drought stress at later stages of stamen maturation (Figure 2D). However, the frequencies of the different (de)hydration types show some variability between different experiments, so that we cannot conclude that the initial stage of a drought-stressed inflorescence unequivocally determines the final hydration status.

**Figure 2.**
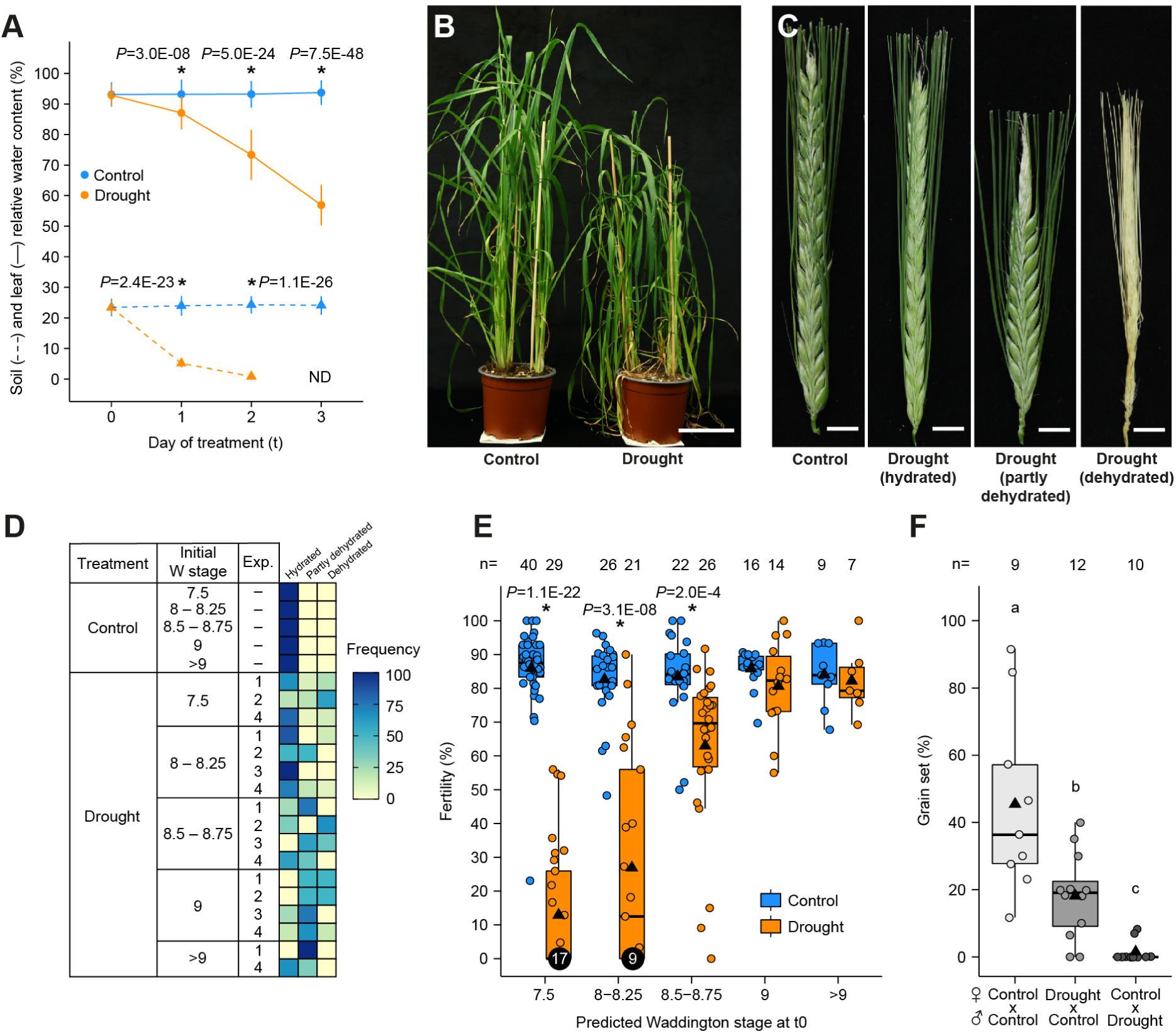
Effects of drought stress applied during Scarlett’s stamen maturation phase. **A**) Soil water content (dash lines) and relative water content of the first tiller’s penultimate leaf (solid lines) under control and drought stress conditions during a 3-day period. Data pooled from three independent experiments (n= 48 [leaf relative water content] and 24 [soil water content]) and plotted as mean ±SD. Asterisks indicate significant differences between control and drought at each time point (one-tailed t-test). Soil water content at day 2 was detectable in only 15 out of 24 drought samples. ND, not detectable. **B**) Representative plant morphology three days after the start of a drought trial. Bar, 5 cm. **C)** Representative spike morphologies around two weeks after the end of a drought trial. Bars, 1 cm. **D)** Frequency of spike hydration phenotypes around two weeks after the end of four independent experiments (Exp.). Frequencies were calculated within all inflorescences predicted to be at the same W stage at the start of the drought stress, and are presented separately for each experiment. Stages W7 – W7.5 and > W9 were not found in every experiment. **E**) Final fertility of hydrated spikelets after the drought treatments start (t0) at different maturation W stages. Fertility calculated as the ratio of filled spikelets over the total number of spikelets. Box plot whiskers represent ±1.5x the interquartile range; triangles, means; horizontal lines, medians; circles, individual measurements. Data pooled from four independent experiments. Numbers above the boxes correspond to sample sizes (n) for each group. Asterisks indicate significant differences between control and drought at each stage, determined with one-tailed t-tests. **F**) Grain set in cross-pollination tests, calculated as the number of grain-filled spikelets over the total number of crossed spikelets. Letters indicate significant differences between the crosses determined with a Conover-Iman test after a Kruskal-Wallis ANOVA (*P* < 0.05). Box plot features as in (E).

While drought seems to elicit an overall fatal stress in fully dehydrated spikelets, we hypothesize that it specifically impairs the development of floral organs in hydrated-but-sterile spikelets. Therefore, we quantified the impact of drought on the fertility of hydrated spikelets. When the treatment started at stages W7.5 and W8–W8.25, either inflorescences were completely sterile (58% and 42% of all spikes analyzed, respectively) or their fertility dropped an average of 73% and 55%, respectively (Figure 2E). Spikelet fertility declined an average of 20% when the treatment started at W8.5–W8.75, while it remained unaffected when started at stage W9 or later (Figure 2E). These findings suggest that inflorescences at the early phase of stamen maturation undergo drought-sensitive processes that are essential for later fertility, whereas inflorescences at later stages are more robust to drought stress, as long as they do not reach a critical dehydration point. We selected stages W8–W8.25 as starting point of all subsequent drought experiments of this work, to specifically target the stamen maturation phase and avoid spikelets that had not exited meiosis.

To clarify if drought impairs the development of male and/or female reproductive organs, we conducted reciprocal crosses between control and drought-treated plants. The efficiency of hand-pollinated control crosses was variable and only achieved an average grain setting rate of 45% (Figure 2F). Yet, when we pollinated pistils of drought-treated plants with fresh control pollen, grain set was significantly reduced to 18% on average (Figure 2F). Moreover, most spikes of control plants that were pollinated with drought-stressed pollen showed no grains (Figure 2F). These results indicate that defects in the development or function of both male and female floral organs can contribute to the loss of fertility induced by drought from the early phase of barley stamen maturation. Nevertheless, male fertility defects alone may explain most of the reduced grain production under drought.

We also followed the progress of stamen and pistil morphology in a 6-day time course during and after the drought treatment. Floral organs of control florets advanced approximately by one to two Waddington stages per day (Figure 3A). During the first three days of treatment, the morphology of stamens and pistils in all drought-stressed florets was indistinguishable from the control, and progressed at the same pace to reach stages W8.75 and W9 (Figure 3A). Thereafter, only 28% of the drought-treated florets examined between days 4 and 6 continued this trend, showing a normal morphology and reaching the end of maturation by day 6 (“drought, unaffected stamens”, Figure 3A). Instead, the remaining 72% of the florets in the stress group exhibited an altered stamen morphology (“drought, affected stamens”, Figure 3A). Still, regardless of the stamen morphology, we did not observe a slowdown in floret development according to the Waddington scale, which is mainly based on pistil features. By contrast, short-term drought stress causes visible changes in plant morphology (Figure 1A). Hence, we matched the Waddington stages of the inflorescences collected during the 6-day time course to the flag leaf sheath extension and the auricle-to-spike-tip distance of their tillers. These two length features were significantly reduced in drought-treated tillers from stages W8.75 and W8.5 onwards, respectively (Figure 3B). Since these stages occurred mainly during the first days of the time course, these results show that drought starts impairing vegetative growth of tiller tissues during the stress treatment, whereas drought-induced changes in floret and stamen morphology occur later, after the end of the stress period.

**Figure 3.**
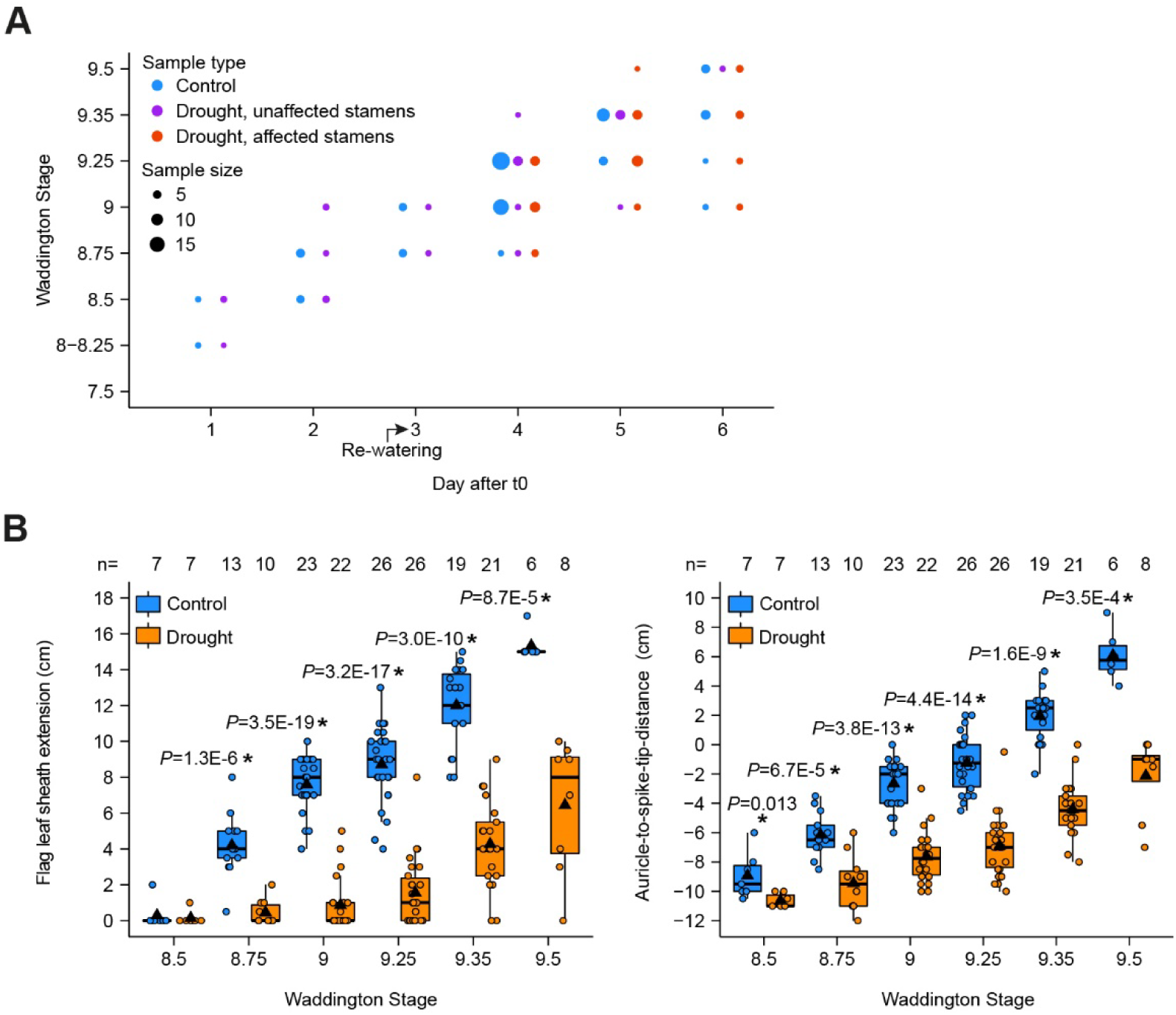
Progression of Scarlett reproductive and vegetative development under drought. **A**) Time course of stamen maturation stages according to our modified Waddington scale. **B**) Flag leaf sheath extension and auricle-to-spike-tip position of tillers with inflorescences at different Waddington stages. Box plot features as in Fig. 2E. Asterisks indicate significant differences between control and drought at each stage (one-tailed t-test). All data in this figure originates from 10 independent experiments.

### Short-term drought during stamen maturation blocks pollen starch accumulation

We then focused on identifying the specific defects that drought induces in stamen maturation and that ultimately account for the loss in male fertility. First, we investigated pollen starch accumulation, an essential feature of cereal crops that is required for pollen fertility and that occurs at late stamen maturation stages in barley (Amanda et al., 2022). Under control conditions, starch becomes visible as individual granules in Scarlett pollen at stage W9.35 and fills the entire pollen grain at stage W9.5. Concomitantly, anthers increase in size and change color from green to yellow (Figure 4). Drought-treated stamens that maintain a mostly normal morphology also accumulate pollen starch between stages W9.35 and W9.5 (Figure 4, unaffected). In contrast, drought-affected anthers not only appear pale yellow and smaller from stage W9.25 but also display no starch at any maturation stage (Figure 4, affected). From here on, we used this obvious change in stamen morphology as indicator of failed pollen starch accumulation caused by drought stress. However, we have not formally tested if pollen grains from drought-unaffected stamens are fertile. Even if they appear to accumulate starch normally, they could be impaired in other functions necessary for fertility, such as pollen tube germination or growth.

**Figure 4.**
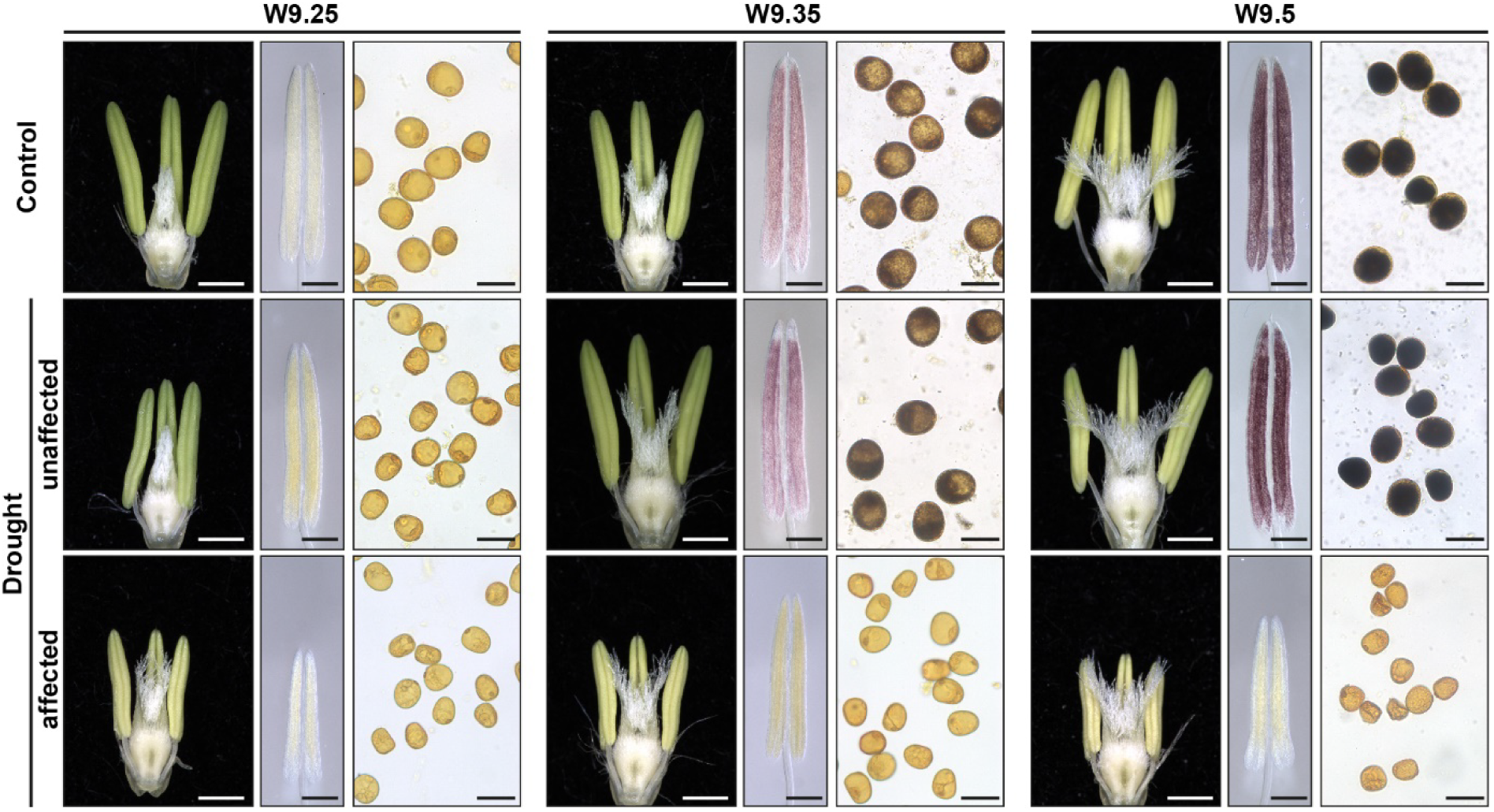
Stamen morphology and pollen starch filling during late maturation stages. Each panel displays reproductive organs (left, bars: 1 cm) and potassium iodide staining in whole anthers (middle, bars: 500 µm) or in pollen grains (right, bars: 50 µm). Iodide reveals starch deposition with a purple or dark blue color.

Second, we evaluated the products of pollen mitosis with DAPI staining of pollen nuclei. This revealed that both control and drought-affected pollen contain a vegetative nucleus and two sperm cells at stage W9.5 (Figure 5A), indicating that pollen mitosis occurred under drought. Third, we quantified pollen viability with a double staining of fluorescein diacetate (FDA) and propidium iodide (PI), which distinguish alive and dead cells, respectively (Figure 5B). We found that, in average, 86% of control and 78% of drought-affected pollen are viable, with only a slight difference between the treatments (*P =* 0.053, figure 5C). Together, the results support that the majority of pollen grains survive the stress and are able to complete two rounds of mitosis. However, drought impairs pollen starch accumulation in 72% spikelets and we propose that this is likely a major cause of reduced male fertility and grain set after water deficiency.

**Figure 5.**
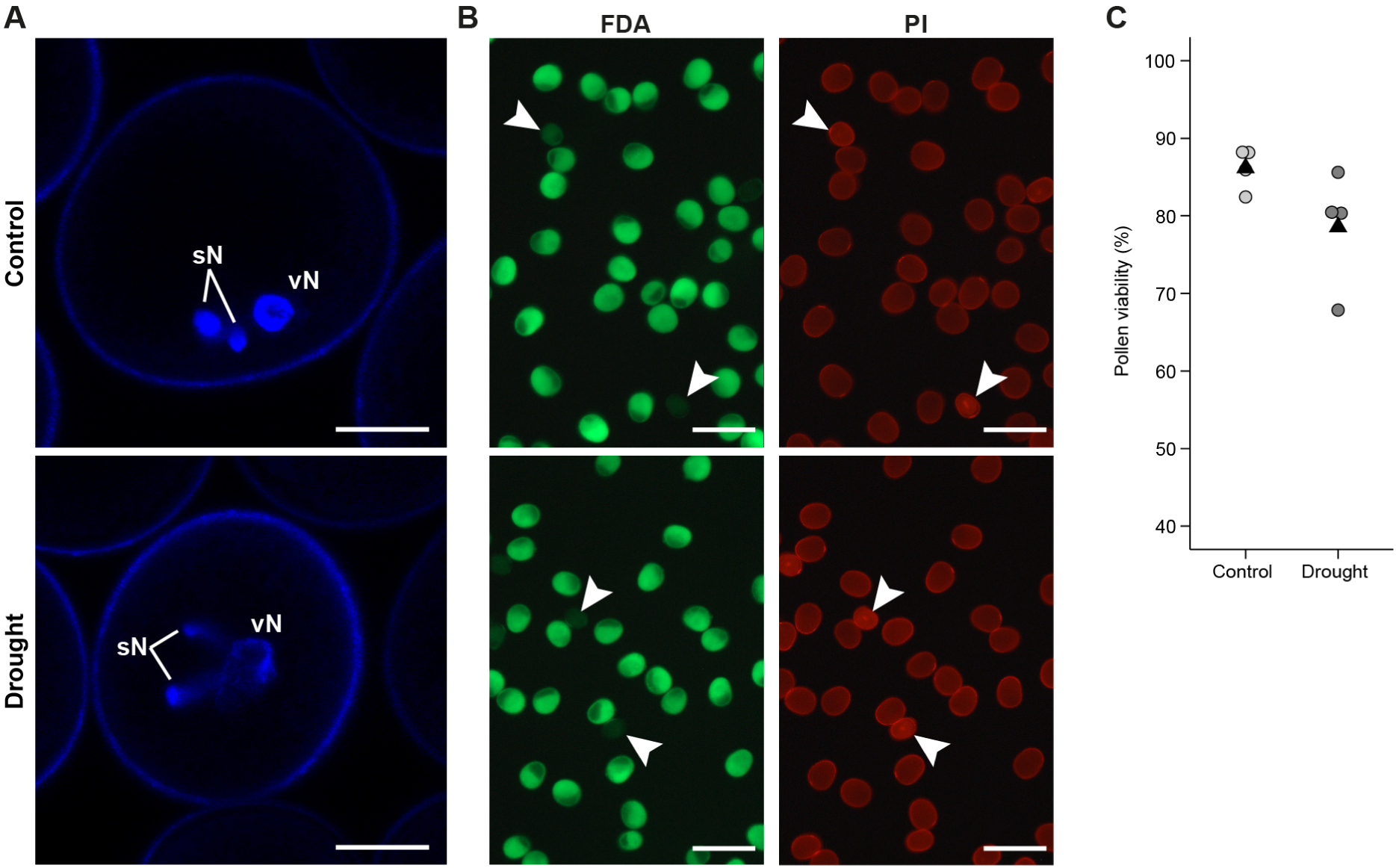
Characterization of pollen at stage W9.5. **A**) DAPI staining of pollen nuclei. vN, vegetative nucleus. sN, sperm cell nuclei. Bars, 10 µm. **B**) Pollen viability in control or drought-affected stamens (confirmed failure of starch accumulation) assayed simultaneously with fluorescein diacetate (FDA) and propidium iodide (PI) dyes. FDA fluoresces (green) in viable cells. Arrowheads indicate pollen with only background fluorescent signal, and thus considered non-viable or dead. Instead, PI accumulates inside dead cells and increases the intensity of fluorescence (red, arrowheads). Note that all pollen grains have a relatively high red signal, a combination of both autofluorescence and emission from PI accumulated on the surface of viable cells (images obtained with a normal epifluorescence microscope). Bars, 200µm. **C**) Quantification of pollen viability determined by FDA-PI staining. Each data point is the ratio of viable pollen counted from three different florets in one inflorescence. Significance of difference between control and drought was determined with a one-tailed t-test. n = 4 inflorescences (598 and 464 total pollen grains scored for control and drought, respectively).

### Short-term drought affects auxin signaling and energy metabolism in maturing stamens

To study the consequences of drought in gene expression, we analyzed the transcriptome of Scarlett stamens at stages W8.5 and W8.75, collected on days two and three of the treatment, respectively. Principal component analysis of all gene expression data reveals that the differences between the transcriptomes of W8.75 stamens under drought and all other samples explain 51% of the total variance (Figure 6A). The second component separates control stamens at W8.75 from the other samples and explains 27% of the variance, probably reflecting the developmental differences between the two stages analyzed. Sample distance analysis confirms this overall separation of the transcriptomes (Figure 6B). We identified 5,598 differentially expressed genes [DEGs, |fold change (FC)| ≥ 1.5; FDR-adjusted P ≤ 0.05] between control and drought samples. The number of DEGs increases from 1,008 at stage W8.5 to 5,183 at W8.75. Altogether, the data indicates that by the third and final day of drought, when the relative water content in vegetative tissues reaches a minimum, the transcriptome of stressed maturing stamens undergoes a major reprogramming.

**Figure 6.**
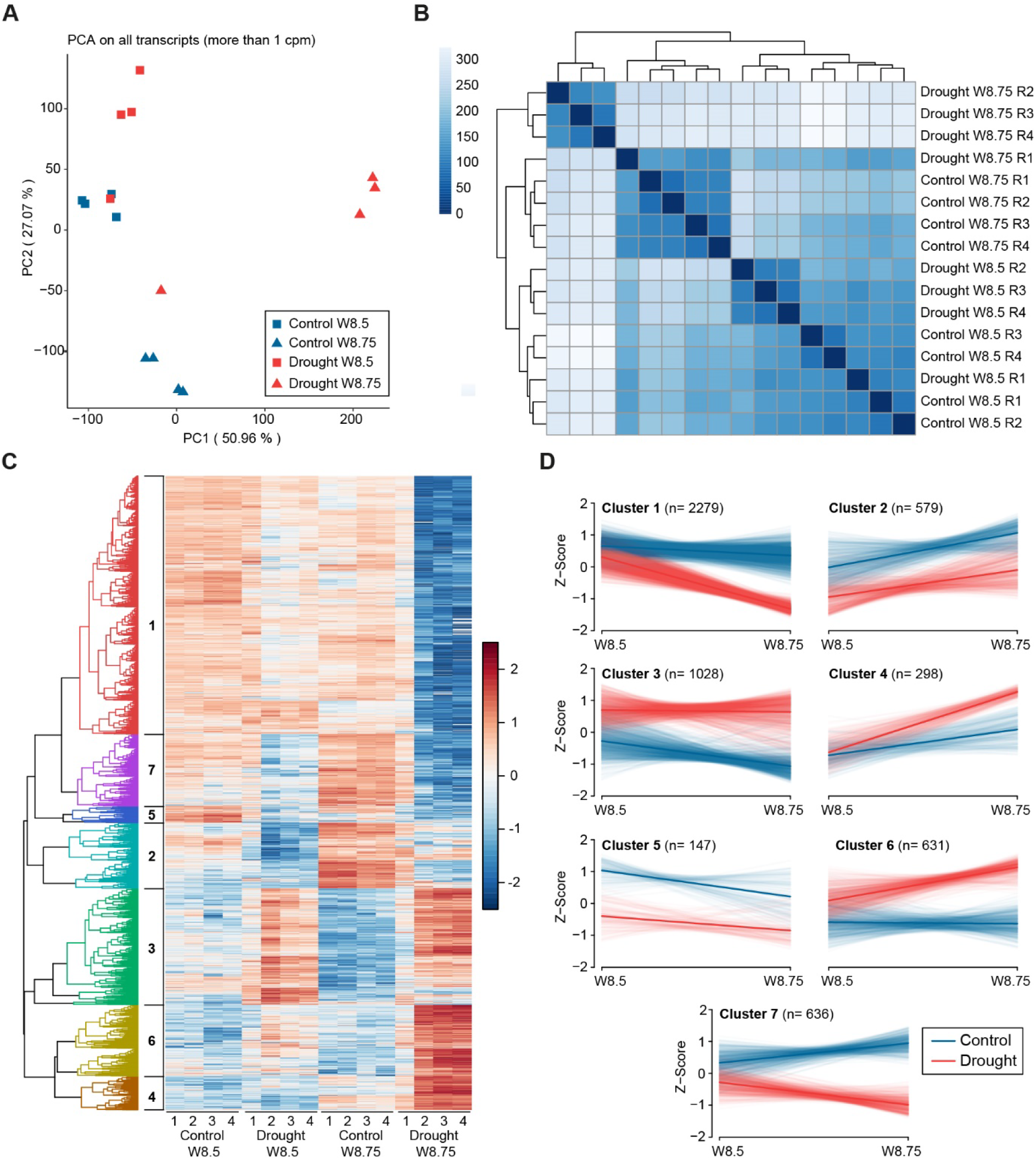
Transcriptomic analysis of Scarlett stamens under drought. **A**) Principal component analysis of normalized expression levels (counts per million, cpm) of all expressed genes. **B**) Heatmap of Euclidean distances between samples, calculated from normalized cpm values of all expressed genes. **C**) Hierarchical clustering of DEGs. Each column is a biological replicate (n = 4 per treatment and stage). Scale represents mean-centered cpm values. **D**) Expression profiles of all DEGs within each cluster. Z-scores are mean-centered and scaled transcript levels in cpm. Light-colored thin lines connect Z-scores at each stage of individual transcripts. Dark-colored thick lines connect the mean Z-Scores across all transcripts at each stage.

DEGs form seven clusters, four of which (1, 2, 5 and 7) contain mostly drought-downregulated genes (65% of all DEGs), while the remaining clusters harbor drought-upregulated genes (Figure 6C, D). To investigate the putative role of DEGs, we conducted a gene ontology analysis on each cluster. Upregulated clusters are enriched for genes related to water and osmotic stress responses, proline synthesis, trehalose metabolism, and signaling through stress hormones (jasmonate, ethylene, ABA). These upregulated pathways may represent stamens’ attempts at resisting the drought stress.

Clusters 1 and 7 contain the genes most strongly downregulated by drought at stage W8.75. These two clusters differ only because genes in cluster 7 seem expressed at lower levels already at stage W8.5, while downregulation of genes in cluster 1 is obvious mostly at W8.75. Both clusters include, among others, genes associated to auxin signaling and carbohydrate metabolism. These categories are in agreement with our recent report that barley pollen produces auxin to boost the expression of energy-generation pathways, which in turn sustain the production of pollen starch (Amanda et al., 2022). In that work, we curated all barley genes associated to auxin signaling, sugar transport and heterotrophic ATP production, and found specific subsets that are putatively induced by auxin and expressed at high levels during the starch filling stages W9.25 and W9.35. Therefore, we followed the expression kinetics of these genes under drought at different stages. In addition to the W8.5 and W8.75 stamen transcriptomes, we obtained additional transcriptomes at W9.25 and W9.35, from stamens with altered morphology collected on days 2 and 3 after the end of the stress. At these later stages, drought affects the expression of 12,188 genes, which represent 62% of the entire transcriptome (19,584 detected transcripts) and a two-fold increase in the number of DEGs from stage W8.75. Plotting the expression kinetics of all DEGs related to auxin signaling, sugar transport and heterotrophic ATP production unveiled three general patterns (Figure 7A). The first and most obvious corresponds to genes strongly downregulated under drought at stages W9.25 and/or W9.35. Every subcategory analyzed contains genes of this type, and several of these genes already show reduced expression at W8.75, particularly in the subcategories “auxin response”, “sucrose hydrolysis” and “glycolysis”. The second pattern contains a few genes downregulated only at stage W8.75, and belonging to the subcategories “auxin response”, “Auxin Response Factors” (ARFs) and “glycolysis”. The third pattern includes genes upregulated under drought, which are also found in almost every subcategory.

**Figure 7.**
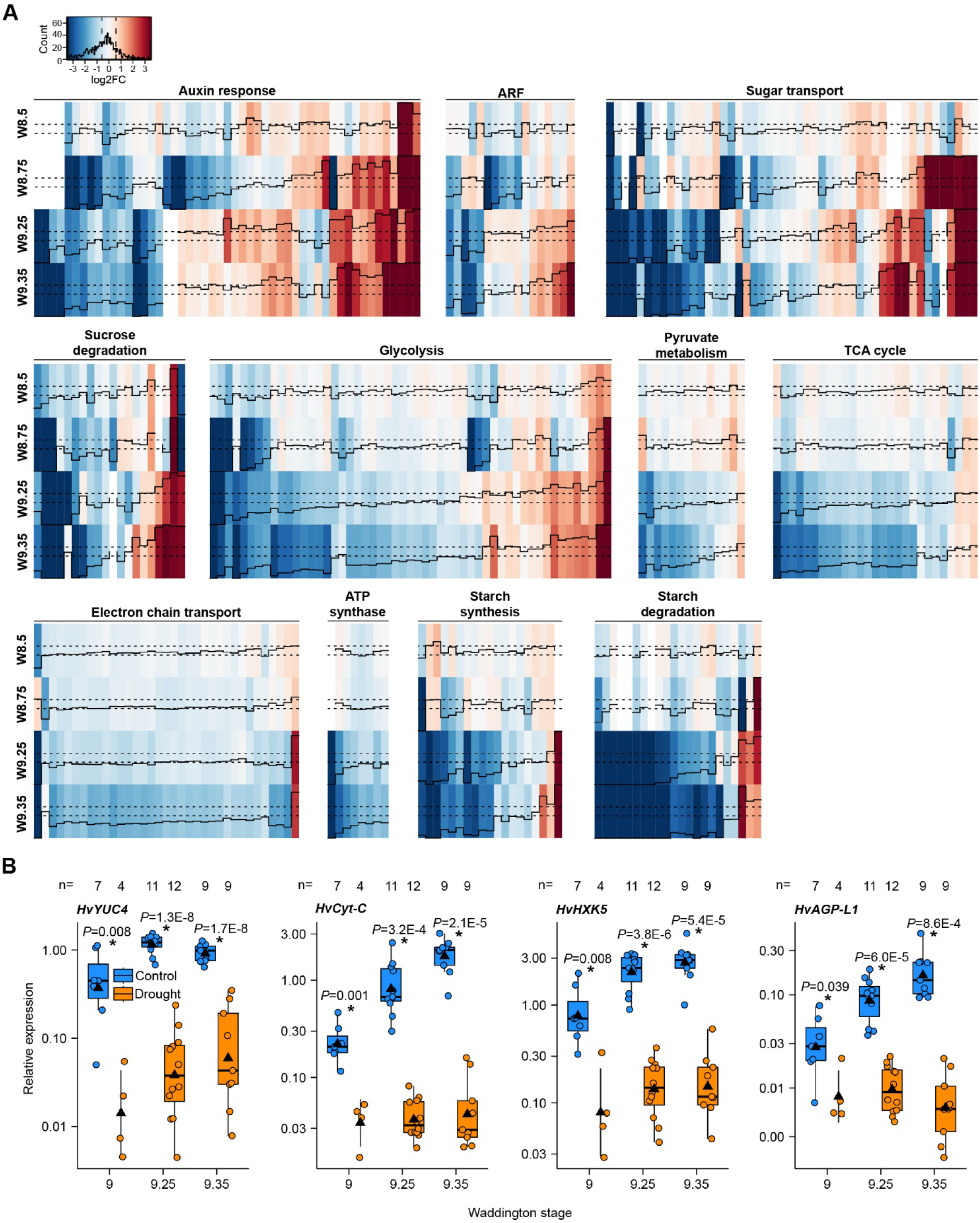
Drought stress during stamen maturation affects the expression of genes related to auxin signaling, sugar transport and energy metabolism. **A**) Heatmaps of log2 fold change values (FC, black solid line) of genes from different subcategories. Each plotted gene is differentially expressed under drought at least at one Waddington stage. Dashed black lines indicate log2 FC values at +/− 0.585 (FC = +/− 1.5), which is our chosen threshold for differential expression. Blank cells without a trace line indicate no detected expression at that particular stage. **B**) qRT-PCR of representative DEGs. Box plot features and statistics as in Fig. 2E, except for drought stage W9, which shows only the mean, SD and the individual data points of the four biological replicates. Total number of biological replicates per category (n) is a pool from two independent experiments.

Barley pollen synthesizes bioactive auxin through the flavin monoxygenase HvYUC4 and the expression of the *HvYUC4* gene increases exponentially during pollen maturation (Amanda et al., 2022). Thus, we hypothesized that the lower expression of auxin signaling genes under drought may be associated to reduced levels of *HvYUC4* transcripts. Indeed, our transcriptome data detected a 13-fold downregulation of this gene under drought at stages W9.25 and W9.35, although its expression appears normal at stages W8.5 and W8.75. qRT-PCR in samples collected independently at stages W9, W9.25 and W9.35 confirmed the downregulation of *HvYUC4*, two energy production genes (*HvHXK5*, glycolysis; *HvCyt-C*, electron transport chain) and the starch synthesis gene *HvAGP-L1* (Figure 7B). In sum, our analyses support that drought may impair pollen starch accumulation by blocking normal expression of genes required for auxin synthesis and signaling, sugar transport and utilization, heterotrophic energy production, and starch synthesis.

To test this interpretation, we evaluated the dynamics of sugars, pyruvate and TCA cycle organic acids in maturing stamens and assessed whether drought affects the levels of those metabolites at the starch-filling stage W9.35. Under control conditions, the contents of sucrose, glucose, fructose and trehalose decline after stage W9.25 (Figure 8A). This correlates with an increase in succinate, the entry point from the TCA cycle into the mitochondrial electron transport chain, at W9.35 and W9.5. On the other hand, pyruvate, citrate and malate tend to decline from stages W9-W9.25, similar to the sugars, while oxoglutarate remains stable (Figure 8A). However, citrate levels increase sharply at final stage W9.5. The overall kinetics supports that starch accumulation in pollen grains normally correlates with increased sugar utilization and channeling of TCA cycle intermediates into succinate likely to support energy production.

**Figure 8.**
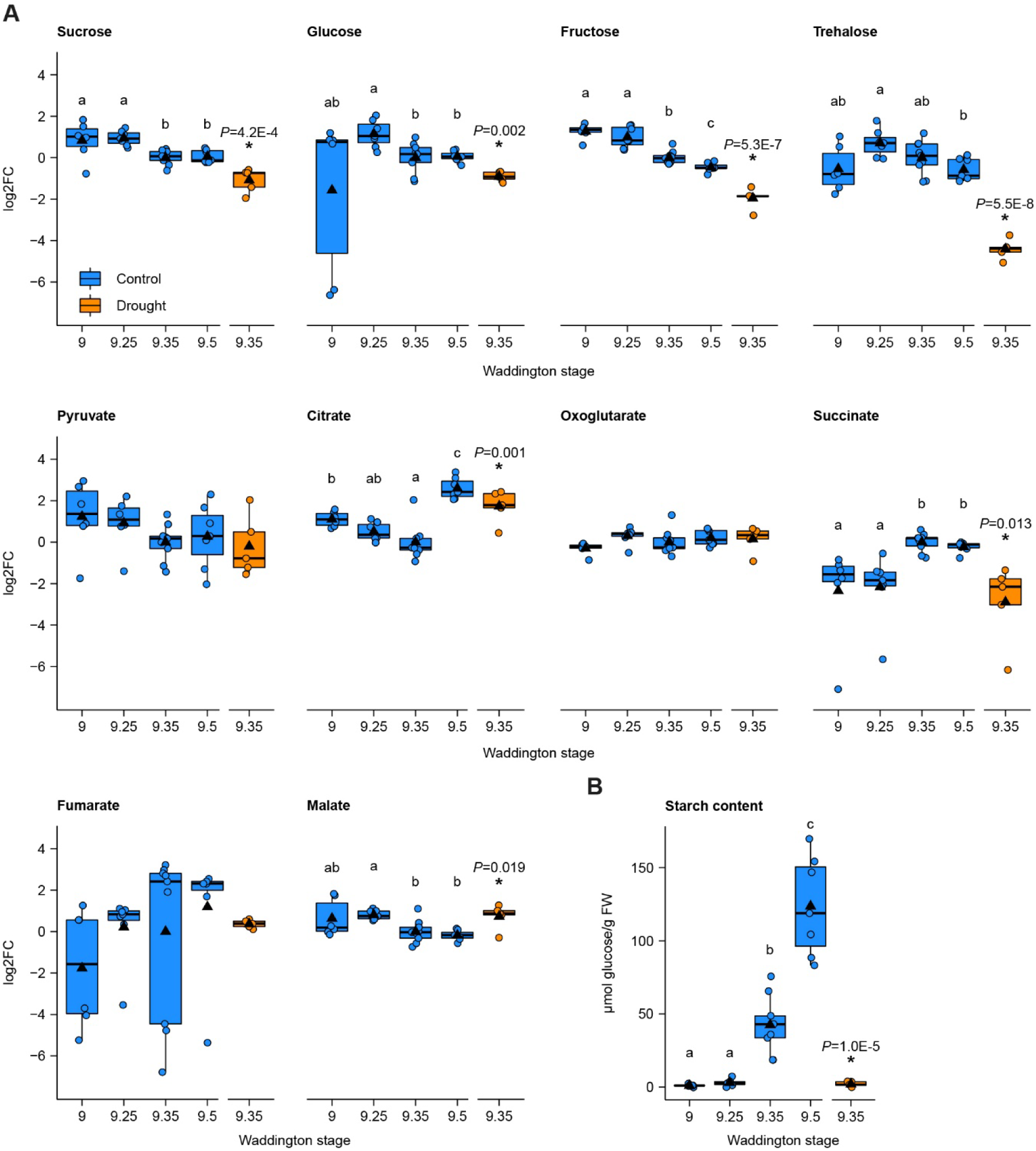
Metabolite profiling of stamens during maturation. **A**) log2 relative levels of disaccharides, hexoses, pyruvate and TCA cycle organic acids calculated from the ratio of normalized values of each replicate to the mean of control W9.35 values. **B**) Starch content. Box plot features as in Fig. 2E. Letters indicate significant differences between control stages determined with a Conover-Iman test following a Kruskal-Wallis ANOVA (*P* < 0.05). Asterisks indicate significant differences between control and drought at stage W9.35, determined with a one-tailed t-test. n= 6 (control W9); 7 (control W9.25, W9.5); 9 (control W9.35); and 5 (drought W9.35), except for pyruvate control W9.25 (n=6), where one outlier with the value 4.62 was excluded from the analysis.

In drought-affected stamens, all sugars are significantly lower at stage W9.35 compared to control stamens at the same stage, and this is also associated to lower succinate levels (Figure 8A). However, pyruvate and oxoglutarate remain unchanged during the stress, while citrate over accumulates (Figure 8A). We confirmed that under control conditions starch levels of Scarlet stamens at stages W9 and W9.25 are low, then increase exponentially at W9.35 and W9.5 (Figure 8B). In agreement with the starch staining (Figure 4), this increase fails to occur in drought-stressed stamens (Figure 8B). Overall, the results suggest that drought leads to the disruption of photosynthate acquisition by the stamen sink tissue, which in turn reduces the flow of succinate into the electron transport chain for ATP generation. Ultimately, this prevents pollen starch accumulation.

## DISCUSSION

### Effects of drought stress on barley stamen maturation

In this work, we established a method to apply a short-term water deficiency stress during the phase of stamen maturation in barley cultivar Scarlett. This simulated drought episode strongly reduces the leaf relative water content and visibly stunts vegetative growth, yet plants survive the stress and continue their life-cycle. Still, in some cases, the stress leads to full dehydration of individual spikelets or complete inflorescences, suggesting that drought may be fatal for spikelet tissues when it exceeds a critical level. Additionally, we found that initiating drought stress during the early stamen maturation phase causes a dramatic reduction of spikelet fertility, while no or only slight effects are seen when the stress starts at advanced maturation stages. Wheat inflorescences also show a stage-specific response to short-term water deficiency (Ji et al., 2010).

Our reciprocal crosses indicate that defects in both male and female reproductive structures may cause the reduction in spikelet fertility under drought, although male sterility is seemingly the major contributor. This agrees with the accepted view that development of male organs in cereals is the reproductive process most vulnerable to drought stress (Dolferus et al., 2011; Yu et al., 2019). Moreover, several studies have documented that drought does not affect ovule fertility (Bingham, 1966; Saini and Aspinall, 1981; Ji et al., 2010). Nevertheless, pollination experiments with drought-stressed pistils have shown significant reduction in grain set of four wheat genotypes (Onyemaobi et al., 2017). Future work should examine drought-treated Scarlett pistils in more detail throughout maturation to identify potential abnormalities, for example in female gametophyte development or function. Moreover, assays investigating stigma receptivity could test if drought disturbs the interaction between pollen and pistils. Further research can also use our short-term drought method to investigate if the sensitivity of stamens and ovaries to water deficiency differs between multiple barley accessions. The natural genetic variation underlying such potential differences could then be used to identify loci for drought tolerance in different reproductive tissues.

We also found that water deficiency does not affect the viability nor the completion of two mitosis rounds in the pollen of cultivar Scarlett. However, it prevents pollen starch accumulation, which is likely the most important cause of the male sterility that we observed under drought. A failure to accumulate starch in pollen has also been associated not only to drought but also to cold or heat stresses in other cereals (Oliver et al., 2005; Ji et al., 2010; Liu and Bennett, 2011; Jin et al., 2013; Begcy et al., 2019). Moreover, Scarlett anthers that harbor starchless pollen are pale and seem smaller than normal, indicating that drought not only affects male gametophyte development but also anther sporophytic tissues.

We did find that a proportion of drought-treated stamens maintain a normal morphology and produce pollen that accumulate starch. We hypothesize that this is the pollen that contributes to the partial and variable grain set that we observed under drought. This should be tested with further pollination experiments with drought-treated stamens bearing either starch-replete or starchless pollen. It is possible that the variable fertility between inflorescences of independent plants reflects differences in water deficiency between pots and plants. However, the partial fertility within inflorescences indicates that some spikelets may withstand the stress better than others. Small variations in developmental timing between spikelets of the same inflorescence could lead to differences in sink capacity, that is, spikelets that are more developmentally advanced might be stronger sinks than their slightly younger counterparts and benefit first from the limited carbohydrate resources under water deficiency. To identify molecular mechanisms or processes of stamen maturation that favor pollen starch accumulation in some spikelets, it would be interesting to compare transcriptomes of drought-affected versus drought-unaffected stamens. The transcriptome replicates of early stages W8.5 and W8.75 included each one drought-treated replicate with transcriptional profiles intermediate between the control and the rest of the drought samples (Figure 6 A – C). We hypothesize that such replicates contained stamens less affected by the water stress, which then could go on to have normal morphology and starch accumulation at later stages.

### Impact of drought on stamen carbohydrate metabolism and energy generation

Previous work has presented limited analyses of gene expression and/or metabolite changes under water deficiency stress in cereal reproductive tissues (Dorion et al., 1996; Ji et al., 2005; Ji et al., 2010; Jin et al., 2013; Li et al., 2015). Our large-scale transcriptome and metabolite analyses in staged barley stamens provide a wider picture that supports a more accurate and plausible model of the negative molecular effects of short-term drought on maturing cereal stamens. In this model, drought directly impairs photoassimilate supply to barley stamens, which deprives pollen of the carbon it requires for starch synthesis. Moreover, we propose that another contributing factor is the strong downregulation of the pollen transcriptional network of auxin signalling and central carbon metabolism genes that is essential for pollen starch production and that depends on the auxin synthesis gene *HvYUC4* (Amanda et al., 2022). The correlated decrease of sugars and *HvYUC4* transcripts after drought stress supports not only such an interpretation but also our recent hypothesis that the build-up of sugars in stamens or pollen may signal the initiation of pollen auxin synthesis (Amanda et al., 2022). In this form, the model suggests that the primary effect of drought is carbon deprivation, while the block of auxin signalling and downstream pathways would be a more downstream, indirect consequence. However, the model also implies a regulatory role of auxin in activating heterotrophic ATP generation pathways not only at the correct pollen development stage but also when resources permit.

Both drought and loss of *HvYUC4* result in reduced expression of similar genes participating in auxin signalling, sugar utilization and ATP synthesis pathways. For example, 115 out of 198 drought-responsive genes related to central carbon metabolism were reported as auxin-dependent by Amanda et al. (2022). This suggests that drought partially mimics the physiological and molecular effects of impaired auxin synthesis. However, one important difference is that drought additionally upregulates a significant number of alternative paralog genes in these same pathways. We hypothesize that this is an attempt of stamen tissues to counteract the negative effects of drought.

Loss of HvYUC4 function in the male sterile mutant *msg38* causes downregulation of the *HvYUC4* gene in stamens as early as W8.25 – W8.5, which correlates with low auxin synthesis at these stages (Amanda et al., 2022). In contrast, our transcriptome and qRT-PCR data indicate that *HvYUC4* expression is affected only after the end of short-term drought, from stage W9. Thus, we hypothesize that auxin synthesis may occur normally during the water deficiency treatment but it starts to fail once the soil and plant water content reach a minimum threshold and does not recover even after plants are re-watered. This idea would also suggest that water deficiency might exert a more direct effect on pollen auxin synthesis. It will be important to quantify auxin levels of drought-treated barley stamens to test these hypotheses.

It is likely that drought hampers leaf photosynthesis, causing decreased photoassimilate production and altered transport of sugars to sink tissues (Barnabas et al., 2008; Farooq et al., 2009; Basu et al., 2016; Du et al., 2020). Accordingly, our stamen transcriptome analysis shows that the expression of genes encoding sugar transporters is extensively altered under drought stress. Moreover, we found expression changes in the barley ortholog of the sugar partitioning regulator *Carbon Starved Anther* (*CSA*), an R2R3 MYB transcription factor that is essential for rice male fertility (Zhang et al., 2010). Reduced expression of sugar transporters may be a direct consequence of lower photosynthate production in source tissues. This is in contrast to loss of *HvYUC4* in *msg38*, where only a relatively small number of sugar transporters is affected (Amanda et al., 2022).

Another major difference between auxin deficiency and drought in barley stamens is the effect in the levels of small organic acids of the TCA cycle. Auxin loss results in a clear decrease of pyruvate, citrate and succinate, all major intermediates of the ATP generation machinery (Amanda et al., 2022). However, drought only reduces the amount of succinate, the entry point to the electron transport chain (Fernie et al., 2004; Schertl and Braun, 2014), which reaches highest levels in control barley stamens at starch-filling stages W9.25 and W9.35. Lower succinate levels under drought may suffice to disrupt the flux of the electron transport chain, leading to reduced ATP generation and consequently hampered pollen starch synthesis. It will be interesting to investigate if low succinate levels are a hallmark of drought in the stamens of other cereals, which could be used as a metabolic readout of water deficiency.

On the other hand, pyruvate levels remain unaffected under drought, perhaps as a result of increased expression of alternative genes, decreased conversion of pyruvate into downstream metabolites or enhanced pyruvate formation through alternative sources (e.g. alanine) (O’Leary, 2021). Moreover, citrate and malate increase in drought-affected stamens, an outcome also observed in leaf and fruit tissues of numerous plant species when exposed to water deficit (Timpa et al., 1986; Guicherd et al., 1997; Sağlam et al., 2010; Levi et al., 2011; Obata et al., 2015; Nahar and Ullah, 2018). It is believed that these small organic acids may act as osmoprotectants or antioxidants to alleviate drought-induced damage. Remarkably, we found that citrate levels also increase strongly in mature control stamens at the late stage W9.5 (Figure 8A), when starch-replete pollen seemingly reaches a mostly dehydrated state that precedes anther opening (Amanda et al., 2022). This raises the intriguing possibility that citrate is a general protective molecule to mitigate low water content in barley pollen or anthers, perhaps at the expense of its function in the TCA cycle.

We conducted the metabolite profiling of drought-stressed stamens only at stage W9.35, a starch-filling stage in cultivar Scarlett. However, our transcriptomic data indicates that drought alters stamen processes related to energy and starch synthesis from stage W8.75. Therefore, further metabolite analyses in drought-stressed stamens at all other development stages considered here (W8.5 to W9.25) should determine how early drought stress alters the metabolism of barley stamens.

ABA is one of the most conspicuous hormones in plant abiotic stress responses. It accumulates in reproductive tissues under drought and has been associated with reduced male fertility (Morgan, 1980; Liu and Bennett, 2011; Dong et al., 2017). In wheat inflorescences, this negative effect has been linked to repression of a sucrose invertase important for sugar utilization, while drought-tolerant lines display lower ABA levels than sensitive lines (Ji et al., 2011). Accordingly, our transcriptome analysis shows that drought both induces the expression of ABA synthesis genes (two *NCED*s) and reduces the expression of ABA catabolism genes (two ABA hydroxylases). Thus, it is possible that ABA also contributes to the downregulation of central carbon metabolism in stamens of barley cultivar Scarlett, which ultimately blocks pollen starch accumulation.

Exploiting exotic germplasms is key to identify genetic resources that confer tolerance against abiotic stresses such as drought (Zamir, 2001; Honsdorf et al., 2017). In wheat and rice, studies have already successfully identified tolerant germplasm by applying transient drought during stages shortly before anthesis (Liu et al., 2006; Ji et al., 2010). Our method to apply a short-term water deficiency stress during the phase of stamen maturation should also provide a reliable avenue to identify sources of drought tolerance in barley stamens. Furthermore, our molecular model of the effects of drought on barley stamen maturation is an important framework to understand tolerance mechanisms in such germplasm.

## MATERIALS AND METHODS

### Growth conditions and drought treatments

We first obtained grains of cultivar Scarlett from Ali Naz (currently at Hochschule Osnabrück). All plants were grown in a controlled greenhouse chamber (16 h light, 22°C; 8 h dark 18°C). Grains were sown in 96-well trays filled with a 1:1 mix of soil ED 73 Einheitserde® (Einheitserdewerke Werkverband e.V., Sinntal-Altengronau, Germany) to BVB Substrates (1:1 vermiculite to coconut fiber). After one week, pairs of seedlings were transplanted to round 1-liter pots (15 cm ⌀, 11.5 cm high), filled with 500 g of the same substrate. We used a FOM2/mts field-operated meter (E-Test, Lublin, Poland) to monitor the volumetric soil water content in pots. Measurements were carried out every morning from Monday to Friday, shortly before watering. Plants were watered on demand to maintain the soil water content close to 25%, which we pre-determined as ideal to keep leaf water content above 90% and allow normal plant development and fertility. Moreover, we fertilized plants once per week with 0.25% WUXAL® Super 8+8+6 (Hauert MANNA Düngerwerke GmbH, Nürnberg, Germany).

Drought treatments were started when the first (hidden) inflorescences approached or reached the stages of stamen maturation. The first three tillers were tagged and pots were randomly assigned to control and treatment blocks. In the drought group, water was withheld for exactly three days, whereas soil water content in control pots was maintained at 25%. Thereafter, drought-treated plants were re-watered and maintained at 25% soil water content until harvest.

### Relative water content measurements

Relative water content (RWC) of the penultimate leaf was used as indicator of the plant water status, and was obtained as described by Liu et al. (2006) with some modifications. Sections of the middle part of the penultimate leaf blade were cut every morning throughout the treatment, weighed and immediately transferred into 2-ml tubes, filled with distilled H_2_O. Tubes were kept for 24 h at 4°C. Then, leaf pieces were blotted dry with tissue paper to remove excess water and turgid weight was determined. Leaf pieces were then transferred into empty 2-ml tubes, dried in an oven at 60°C for 48 h and their dry weight determined. RWC was calculated with the equation RWC = 100% x [fresh weight – dry weight] / [turgid weight – dry weight] from Schonfeld et al. (1988).

### Imaging of plants and reproductive structures

Photographs of whole plants and inflorescences were taken with an EOS 500D camera carrying an EF 16–35 mm F/4 L IS USM lens (Canon, Tokyo, Japan). Florets were carefully dissected using fine forceps under a common stereomicroscope, and reproductive organs were placed on black velvet for imaging with a Carl Zeiss (Oberkochen, Germany) Discovery.V12 stereomicroscope plugged to a Stemi 503 color camera. For visualizing microspores within anthers at stages W7.5 – W8.25, anthers were mounted in perfluoroperhydrophenanthren (Sigma-Aldrich/Merck) and imaged with a Zeiss Axio Scope.A1 light (epifluorescence) microscope combined with an Axiocam 512 color camera. Both microscopes were controlled with Zeiss’ ZEN v2.3 blue edition software. Scale bars were added with ImageJ v1.53 (https://imagej.nih.gov/ij/).

### Reciprocal crossing experiments

To prevent self-pollination, spikelets from the middle of inflorescences and containing W9.35 florets were carefully opened with clean forceps, emasculated and covered with plastic wrap. One or two days later, when pistils showed full spreading of stigmatic branches, we sprinkled them with pollen of recently opened anthers. We then covered the inflorescences with waxed paper bags. At maturity, we calculated grain set in individual inflorescences as the fraction of obtained grains over the number of pollinated pistils.

### Pollen starch staining

Anthers were opened with fine forceps and pollen grains were immersed in I_2_/KI solution (0.3 g/1.5 g per 100 ml H_2_O) on glass slides, before imaging with the light microscope. Whole-anther starch staining was conducted on fresh anthers fixed in Carnoy’s solution (6:3:1:: ethanol: chloroform: glacial acetic acid) for 1 – 2 h. Anthers were transferred to new 1.5-ml tubes filled with staining solution (0.5 g / 2.5 g I_2_/KI in 100 ml 8:2:1:: chloral hydrate: glycerol: water) and rotated for approximately 4 – 6 h. Staining solution was discarded and anthers were cleared two times with fresh chloral hydrate solution for 24 h each, always under rotation. Anther samples were mounted in chloral hydrate on glass slides and imaged with the stereomicroscope.

### Pollen nuclei visualization

Each replicate of 6 – 9 anthers at stage W9.5 was placed in 1.5-ml tubes containing 100 µL of 4’6,-diamidino-2-phenylindole (DAPI) staining solution (1 µg/ml DAPI and 1% Triton-X in 1x PBS, pH 7.4) and 15 µL of 0.01% toluidine blue (to remove excess tissue autofluorescence). Samples were vortexed, briefly spun down and flicked gently. Tubes were covered with aluminum foil and incubated for 1 h at room temperature. The bulk of pollen sedimented after a short centrifugation and was carefully pipetted to a microscope slide and mounted on perhydroperfluorophenanthren. Nuclei were imaged with an SP8 confocal microscope (Leica Microsystems, Wetzlar, Germany) running a 405 nm laser beam and a HyD detector capturing the emitted fluorescence between 410 – 505 nm.

### Pollen viability assay

Each replicate of minimum 9 anthers originating from a single inflorescence was placed in 1.5-ml tubes filled with staining solution (1 ml of 1x PBS with 20% sucrose + 10 µL of 2 mg/mL fluorescein diacetate + 20 µL of 1 mg/ml propidium iodide). Samples were vortexed and incubated in the dark for 5 min at room temperature. After brief centrifugation, the sedimented pollen was pipetted into a fresh 1.5-ml tube and washed twice with 20% sucrose in 1x PBS before mounting on a microscope slide for visualization with the epifluorescence microscope.

### RNA extraction and qRT-PCR

Stamen collection took place as follows: stages W8.5 and W8.75, days two and three of the drought-stress treatment, respectively; stage W9, one day after the end of the stress; and stages W9.25 and W9.35, two to three days after the end of the treatment. Each replicate consisted of stamens dissected from florets collected from individual inflorescences: either 13 – 15 florets for stages W8.5 and W8.75, or 7 – 9 florets for W9 – W9.35. RNA extraction, cDNA synthesis and qRT-PCR reactions and analysis were performed as described (Amanda et al., 2022) with oligonucleotides already reported in that study, as follows:

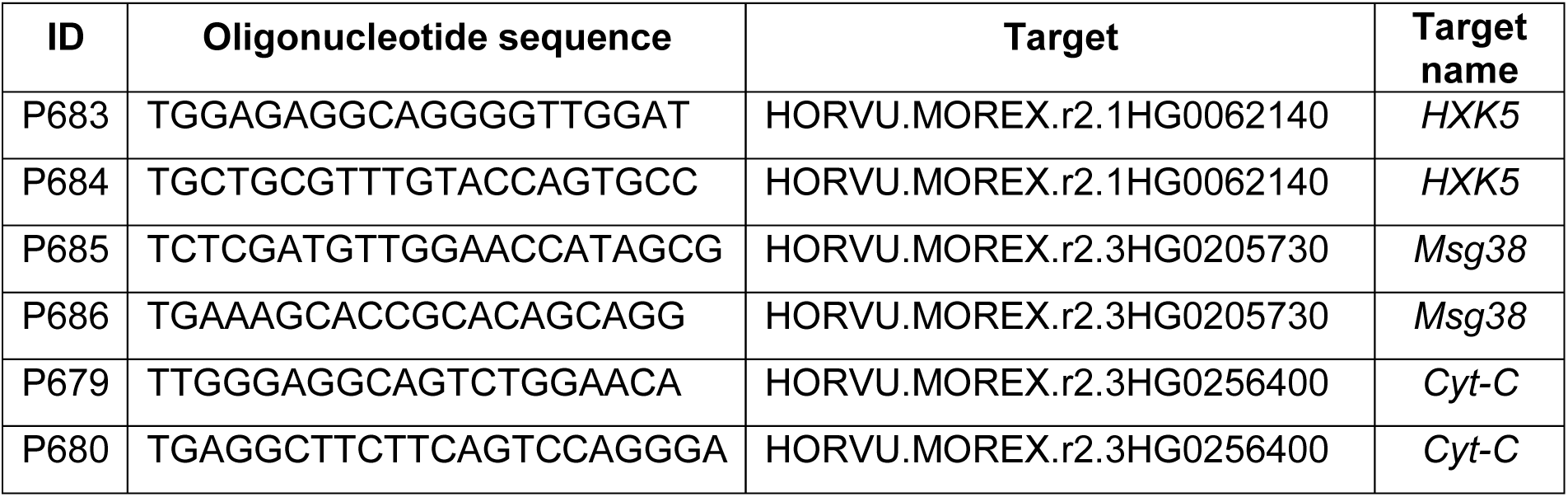
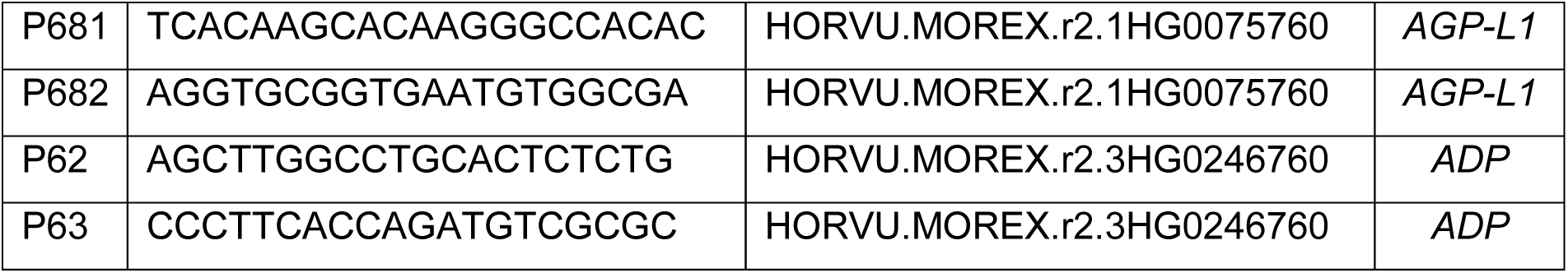

### Transcriptome analysis

We performed two separate transcriptome analyses of Scarlett control and drought-stressed stamens, one with stamens at stages W8.5 and W8.75, the other at stages W9.25 and W9.35. Four biological replicates were collected per stage and treatment. Ribosomal RNA depletion, library preparation and RNA sequencing (20 million 150-bp single reads per sample), was conducted at the Max Planck Genome Centre in Cologne with a HiSeq 2500 System (Illumina, San Diego, USA). All subsequent data analysis was performed as described (Amanda et al., 2022), including quality-check and trimming of sequencing data, transcript quantification with Salmon v1.4 (Patro et al., 2017), data normalization, principal component analysis, Euclidean distance plotting, differentially expressed gene identification, hierarchical clustering and gene ontology enrichment. Moreover, we used the curated list of genes related to auxin signaling, sugar transport and central carbon metabolism reported in that work.

### Metabolite profiling and starch measurements

Sample material was collected in five independent experiments. We dissected stamens from 19 – 34 florets coming from three inflorescences for each W9 – W9.5 control replicate. For each W9.35 drought replicate, we had to pool 29 – 46 florets from four to five inflorescences due to their smaller size. Every replicate contained stamen material originating from the same experiment, except for two W9.35 drought replicates, where stamens from two experiments were pooled. Stamens were collected in 1.5-ml tubes, fresh weight was measured and samples were immediately frozen in liquid nitrogen. In total, 5 – 9 control replicates per stage and 5 W9.35 drought replicates were analyzed. Metabolite profiling and starch measurements were conducted as described (Amanda et al., 2022).

### Statistical analyses

Statistical tests and values of *n* are reported directly in plots or presented in figure legends. Differences of means between control stages were tested in Excel 2019 with one-tailed Student’s t-tests after evaluating homogeneity of variances with *F*-tests. Shapiro-Wilk test was performed in R to test data for normal distribution. Significant differences between means in multiple comparisons were tested by Kruskal-Wallis ANOVA and a subsequent Conover-Iman test with Bonferroni correction. All tests were performed with a significance threshold of P<0.05. Plots were created with R’s package *ggplot2* and compact letter display for Figures 2F and 8 was obtained with the package *multcompView*.

## ACKNOWLEDGEMENTS

This work was supported by the DFG-funded Research Training Group GRK2064: ‘Water use efficiency and drought stress responses: From Arabidopsis to barley’ (I.F.A. via University of Bonn); the Max Planck Society (I.F.A. and A.R.F.); and the European Union’s Horizon 2020 research and innovation programme (grants SGA-CSA 664621 and 739582 under FPA664620 to A.R.F.).

## AUTHOR CONTRIBUTIONS

Conceptualization, I.F.A.; data curation, I.F.A.; formal analysis, R.L. and Y.Z.; funding acquisition, I.F.A. and A.R.F.; investigation, R.L. and Y.Z.; supervision, I.F.A. and A.R.F.; visualization, R.L.; writing—original draft, R.L. and I.F.A.; writing—review & editing, I.F.A. with input from all authors.

